# Gain- and loss-of-function mutations cause perinatal lethality in mouse models of Birk-Barel syndrome

**DOI:** 10.64898/2026.09.08.750008

**Authors:** Sylvain Feliciangeli, Franck C. Chatelain, Frédéric Fiore, Delphine Bichet, Florian Lesage

**Affiliations:** Université Côte d’Azur, CNRS, Inserm, Institut de pharmacologie moléculaire et cellulaire (IPMC), Valbonne, France; Centre d’immunophénomique (CIPHE), Aix Marseille Université, Inserm, CNRS, CELPHEDIA, PHENOMIN, Marseille, France

**Keywords:** K2P channel, genetic syndrome, mouse models

## Abstract

Birk-Barel syndrome (BBS), also known as KCNK9 imprinting syndrome, is a neurodevelopmental disorder caused by mutations in the maternally expressed paternally imprinted KCNK9 gene that encodes the TASK3 potassium channel. The first mutation identified in patients with BBS was a loss-of-function variant that reduces the potassium current mediated by TASK3. Subsequent studies have uncovered additional pathogenic variants, including gain-of-function mutations, that increase TASK3 activity. Here, we show that both mutations resulted in the expression of the hyperactive Task3M159I variant or the hypoactive Task3G236R variant, which caused early postnatal lethality in mice when maternally inherited, confirming the monogenic nature of BBS. These results indicate that the pathogenesis of BBS involves mechanisms other than alterations in channel activity. Consequently, the proposed therapeutic strategies, such as pharmacological modulation of the TASK3 channel function or epigenetic reactivation of the paternal KCNK9 allele, do not seem to be viable options. These findings also demonstrate that TASK3 knockout mice are not suitable models for studying KCNK9 imprinting syndrome or for developing therapies for affected patients. Therefore, there is an urgent need for animal models that allow conditional expression of BBS-associated mutations.

## Introduction

Birk-Barel syndrome (BBS), also known as KCNK9 imprinting syndrome, is a rare disorder (approximately 50 cases have been reported to date) characterized by symptoms of variable severity, including intellectual disability, hyperactivity, epilepsy, congenital hypotonia, and dysmorphic features, most notably an elongated face. Genetic studies have linked pathology to alterations in the KCNK9 gene, which is subjected to paternal imprinting, such that only the maternal allele is transcribed. KCNK9 encodes TASK3, a two-pore-domain potassium channel that is predominantly expressed in the nervous system, where it contributes to the background inhibitory conductance (Feliciangeli et al, 2015). Electrophysiological studies have demonstrated that the first identified mutation, G236R (Barel et al., 2008; Graham et al., 2016), causes a reduction in TASK3 current and renders the channel insensitive to pH and GqPCR-mediated regulation. The activation of residual current using flufenamic acid, an activator of TASK3, has been proposed as a means of alleviating the symptoms of patients (Veale et al., 2014). In a separate study, mice carrying a maternal Kcnk9 knockout allele showed that reexpression of the imprinted paternal Kcnk9 allele through the inhibition of histone deacetylation improved cognition, thereby identifying epigenetic manipulation as a potential therapeutic approach (Cooper et al., 2020).

However, in 2022, Cousin et al. described 15 novel BBS mutations, several of which, including M159I, cause a gain in TASK3 function (Cousin et al., 2022). How both gain and loss of TASK3 function can produce the same syndrome remains unexplained, and this paradox questions the rationale for activating TASK3 or re-expressing the wild-type paternal allele as a treatment strategy. Therefore, animal models are essential for a better understanding of the mechanisms underlying BBS, and would be of considerable assistance in the clinical management of patients. To this end, we developed knock-in mouse models carrying BBS mutations in Kcnk9, resulting in either a loss-of-function mutation (Task3G236R) or a gain-of-function mutation (Task3M159I).

## Material and methods

Electrophysiological experiments were conducted as described previously (Khoubza et al., 2022). Cell surface quantification of TASK3 was performed according to previously established protocols (Oliveira-Mendes et al., 2021). Briefly, a pH-sensitive fluorescent protein was fused to the channel and the constructs were transiently expressed in MDCK cells. The cells were monitored by microscopy while alternating the extracellular medium between neutral and acidic pH conditions. The surface expression ratio was calculated as the percentage of pH-sensitive fluorescence. All animal procedures were carried out in strict accordance with the European Commission Directive 2010/63/EU for the care and use of laboratory animals, and were approved by the French National Ethical Committee (APAFIS#46816). The mouse were used in this study was C57BL/6J. Wild-type (WT) mice were obtained from Janvier Laboratories. Transgenic mouse lines harboring the TASK3 M159I and TASK3 G236R mutations were generated at the Centre d’Immunophénomique (CIPHE) in Marseille. Mice were group-housed (2–5 animals per cage) under standard environmental conditions, including a temperature range of 20–22°C, 40 percent relative humidity, and a 12-hour light/dark cycle (lights off at 20:00). Food and water were provided ad libitum. Genotyping was performed by PCR using tail-tip DNA.

## Results

Fig. 1C shows that BBS mutations introduced in the mouse channel produce the same effects as those observed in the human channel: a marked decrease in Task3 activity with the G236R mutation (1.1 ± 0.2 µA, n = 6, compared to 13.1 ± 2.3 µA, n = 9 at +60 mV) and an increase in activity with the M159I mutation (17.2 ± 1.8 µA at +60 mV, n = 8). These functional changes were not associated with alterations in the channel trafficking to the plasma membrane (Fig. 1B). Using CRISPR-Cas9 technology, two mouse lines were generated, each carrying either the M159I or the G236R mutation in Kcnk9. When heterozygous mutation-bearing males were crossed with wild-type females, the resulting litter contained the expected number of pups. The mutated allele followed Mendelian transmission patterns, with approximately 50% of offspring being heterozygous (Fig. 1C). In contrast, when heterozygous mutation-bearing females were crossed with wild-type males, litter sizes were significantly reduced, and no heterozygous pups were detected at weaning (Fig. 1C). This phenotype was consistent for both mutants, indicating that the maternal transmission of either mutated channel led to early lethality. To assess whether lethality occurred in utero, pregnant heterozygous females were sacrificed at embryonic days 19–20 (E19–E20) and embryos were collected for subsequent analysis. The results present-ed in Figure 1D were consistent across both lines: litter sizes were as expected and transmissions aligned with Mendelian expectations, with approximately 40% of the embryos carrying the mutated Kcnk9 allele. No significant differences in weight or anatomical features were observed between the heterozygous and wild-type littermates. However, the slightly reduced proportion of heterozygous embryos relative to the expected ratio may indicate the potential impact of the mutation during early in utero development. These observations indicated that the absence of mutation-bearing mice at weaning is attributable to postnatal lethality. Pregnant females were subjected to continuous litter monitoring to determine the precise timing of their death. This approach revealed a peak in mortality during the first 24 hours after de-livery. Although it was not possible to recover the bodies of all pups due to maternal elimination, genotyping of the recovered specimens confirmed that they carried the mutated TASK3 gene. Notably, both the constructs yielded comparable results.

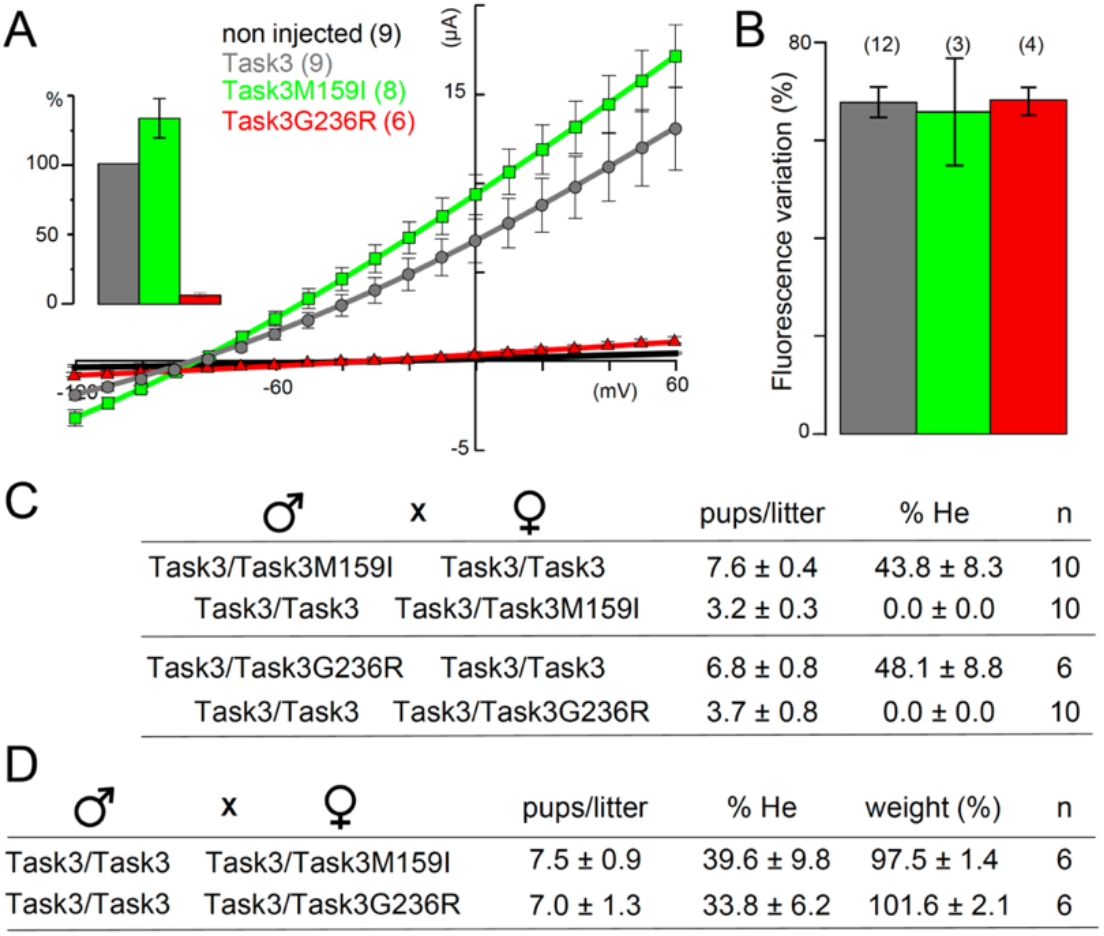
**A.** Electrophysiological recordings of *Xenopus* oocytes expressing Task3 (blue), Task3M159I (green), or Task3G236R (red). Inset: Current amplitude at +60 mV, expressed as a percentage of wild-type Task3 current. **B**. Fraction of channels expressed at the cell surface using the same color coding as in panels A. **C**. Litter sizes and percentage of heterozygous (He) pups at weaning. The breeding combinations are indicated. **D**. Litter sizes and percentages of heterozygous (He) embryos (E19–E20). The breeding combinations are indicated. All values are presented as mean ± standard error of the mean (SEM). *The number of independent experiments (n) is indicated for each dataset*.

## Discussion

These results demonstrated that BBS mutations induce perinatal lethality in mice. Patients with this condition require parenteral nutrition for several months after birth, and exhibit lifelong feeding difficulties. The inability of mutant pups to suckle maternal milk likely underlies their rapid death shortly after birth. Notably, both gainof-function M159I and loss-of-function G286R mutations produce the same phenotype, suggesting that the pathogenesis of BBS involves mechanisms that go beyond simple dysregulation of channel activity.

Furthermore, pharmacological modulation of the TASK3 channel function or epigenetic reactivation of the paternal allele no longer appears to be a viable therapeutic strategy. These findings also underscore that Task3 knockout mice are not an appropriate model for investigating BBS or developing therapies for affected patients. Therefore, the generation of new animal models with conditional gene expression is essential to bypass early postnatal natal lethality and to enable further research.

## Funding Statement

This study was supported by grant SAM-2022-121403 from the Foundation Maladies Rares.

## Data Access Statement

Research data supporting this publication are available from the NN repository upon request.

## Conflict of Interest declaration

The authors declare that they have no affiliations with or involvement in any organization or entity with any financial interest in the subject matter or materials discussed in this manuscript.

## Author Contributions

SF, FCC, FF, and DB designed and implemented the research, SF and FL analyzed the results, and wrote the manuscript. FL conceived and supervised the project.

## Notes

### Competing Interest Statement

The authors have declared no competing interest.

